# *Monodopsis* and *Vischeria* genomes elucidate the biology of eustigmatophyte algae

**DOI:** 10.1101/2021.08.22.457280

**Authors:** Hsiao-Pei Yang, Marius Wenzel, Duncan A. Hauser, Jessica M. Nelson, Xia Xu, Marek Eliáš, Fay-Wei Li

## Abstract

Members of eustigmatophyte algae, especially *Nannochloropsis*, have been tapped for biofuel production owing to their exceptionally high lipid content. While extensive genomic, transcriptomic, and synthetic biology toolkits have been made available for *Nannochloropsis*, very little is known about other eustigmatophytes. Here we present three near-chromosomal and gapless genome assemblies of *Monodopsis* (60 Mb) and *Vischeria* (106 Mb), which are the sister groups to *Nannochloropsis*. These genomes contain unusually high percentages of simple repeats, ranging from 12% to 21% of the total assembly size. Unlike *Nannochloropsis*, LINE repeats are abundant in *Monodopsis* and *Vischeria* and might constitute the centromeric regions. We found that both mevalonate and non-mevalonate pathways for terpenoid biosynthesis are present in *Monodopsis* and *Vischeria*, which is different from *Nannochloropsis* that has only the latter. Our analysis further revealed extensive spliced leader *trans*-splicing in *Monodopsis* and *Vischeria* at 36-61% of genes. Altogether, the high-quality genomes of *Monodopsis* and *Vischeria* not only serve as the much-needed outgroups to advance *Nannochloropsis* research, but also shed new light on the biology and evolution of eustigmatophyte algae.

## Introduction

The diversity of algae is vast but largely unexplored. Despite their often inconspicuous nature, algae have played pivotal roles in Earth’s biogeochemical cycles (De Vargas et al. 2015), and some might hold the key to sustainable bioenergy production (Radakovits et al. 2010; Jagadevan et al. 2018). Eustigmatophytes (Class Eustigmatophyceae) is a small algal lineage in Ochrophyta (Stramenopiles), with around 15 genera and 30 species described to date. They are single-celled coccoid algae that can be found in freshwater, soil, and marine environments. The phylogeny and taxonomy of this group have only been recently clarified (Amaral et al. 2020; Fawley et al. 2014, 2015; Eliáš et al. 2017; Ševčíková et al. 2019).

The eustigmatophytes that have garnered the most attention are undoubtedly *Nannochloropsis* (including the recently segregated *Microchloropsis*; Fawley *et al.*, 2015). Many *Nannochloropsis* species are capable of producing a staggering amount of lipids, up to 60% of the total dry weight. Because of this, as well as their fast growth rate, much research effort has been devoted to developing *Nannochloropsis* as an industrial biofuel alga. The genomes of several *Nannochloropsis* species and strains have been sequenced (Pan et al. 2011; Radakovits et al. 2012; Vieler et al. 2012; Wang et al. 2014; Corteggiani Carpinelli et al. 2014; Schwartz et al. 2018; Ohan et al. 2019; Guo et al. 2019; Brown et al. 2019; Gong et al. 2020), although only a few assemblies have reached high contig continuity and completeness (Fig. 1A). In addition, tools for genetic transformation, gene editing, and marker-less trait-stacking have also been developed (Radakovits et al. 2012; Vieler et al. 2012; Verruto et al. 2018; Poliner et al. 2018; Osorio et al. 2019; Poliner et al. 2020; Wei et al. 2017; Naduthodi et al. 2019). The applications of these tools and resources have resulted in substantial improvements of lipid production in *Nannochloropsis* (Ajjawi et al. 2017).

**Fig 1.**
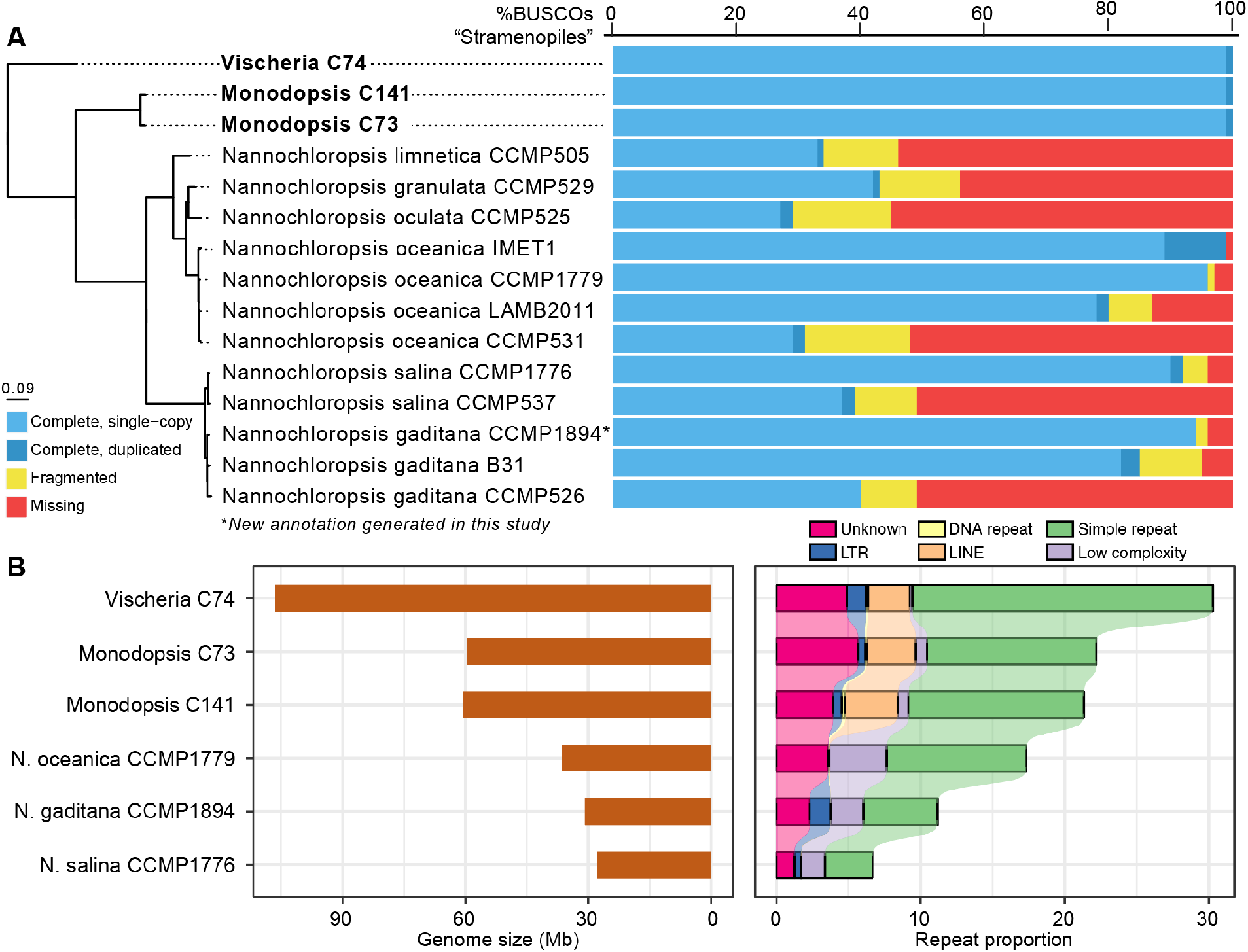
Comparisons of eustigmatophyte genomes. (A) The three genomes reported here (in bold) have the highest BUSCO proteome completeness scores compared to the currently available *Nannochloropsis* genomes. The phylogeny on the left was based on 1302 single-copy loci, and all branches receive bootstrap support of 100. The rooting was determined by OrthoFinder, which is consistent with the published phylogenies (Ševčíková et al. 2019). (B) Overall genome size (left panel) correlates well with repeat content (right panel). Significant expansions of simple repeats and LINE are evident in *Vischeria* and *Monodopsis* genomes.

On the other hand, relatively little is known about the eustigmatophytes beyond *Nannochloropsis*. To date, most of the research on non-*Nannochloropsis* eustigmatophytes has focused on the organellar genomes (Ševčíková et al. 2016; Yurchenko et al. 2016; Huang et al. 2019; Ševčíková et al. 2019) and the association with a novel endosymbiont *Candidatus* Phycorickettsia (Yurchenko et al. 2018). Despite many interesting findings that have recently emerged from non-*Nannochloropsis* eustigmatophytes, the lack of sequenced genomes is limiting further studies.

Here we report three near-chromosomal genome assemblies of *Monodopsis* spp. and *Vischeria* sp., representing two separate lineages that are successively sister to *Nannochloropsis* (Fig. 1A, Fig. S1). We carried out comparative studies of repeats and gene space and found evidence of spliced leader *trans*-splicing in eustigmatophytes. Our results here help to gain a more holistic view on the biology and genomic diversity of eustigmatophytes, expanding beyond what was only known from *Nannochloropsis*.

## Results and Discussions

### Eustigmatophytes isolated from bryophytes

In our ongoing effort to isolate symbiotic cyanobacteria from surface-sterilized bryophyte thalli (Nelson et al. 2019), we have occasionally obtained eustigmatophyte algae instead. DNA barcoding using the 18S rDNA marker indicates all our eustigmatophyte isolates belong to either *Monodopsis* or *Vischeria* (see Fig. S1 for the 18S rDNA phylogeny). So far, we have isolates from multiple species of hornworts, liverworts, and mosses, and from diverse geographic locations spread across North America (Table S1). The nature of interaction between eustigmatophytes and bryophytes (if there is any) is unclear. A symbiotic relationship is a possibility, given that similar algal strains have been repeatedly isolated from bryophytes. The recent finding that *Nannochloropsis oceanica* could enter an endosymbiotic relationship with the fungus *Mortierella* (Du et al. 2019) further speaks to the symbiotic competency of eustigmatophytes. On the other hand, both *Monodopsis* and *Vischeria* are common soil algae, and it is possible that they are resistant to our sterilization method and came out as “contaminants”. Future experiments are needed to examine the possible eustigmatophyte-bryophyte interaction.

### Near-chromosomal level assemblies of Monodopsis and Vischeria

To obtain high quality reference genomes, we generated Illumina short reads and Oxford Nanopore long reads for one *Vischeria* (C74) and two *Monodopsis* strains (C73, C141). The K-mer based genome size estimates were around 60 and 100 Mb for *Monodopsis* and *Vischeria*, respectively. After filtering, the Nanopore data represented 45-67X coverage with a read length N50 between 13-25kb (Table S2). The assemblies based on Flye (Kolmogorov et al. 2019) are near chromosomal, with the majority of the contigs containing at least one telomeric end (Table 1). The telomeric motif is “TTAGGG”, which was also found in *Nannochloropsis* (=*Microchloropsis*) *gaditana* B-31 (Corteggiani Carpinelli et al. 2014). A total of 13,969, 13,933, and 18,346 protein-coding genes were annotated from *Monodopsis* C73, *Monodopsis* C141, and *Vischeria* C74, respectively, all with a 100% BUSCO (Benchmarking Universal Single-Copy Orthologs)(Simão et al. 2015) completeness score against the “Stramenopile” dataset. Compared to the published *Nannochloropsis* genomes, the assemblies we present here are by far the most complete (Fig. 1A). Interestingly, none of the three genomes contain *Ca.* Phycorickettsia contigs that were previously reported in other eustigmatophytes (Yurchenko et al. 2018).

**Table 1.** Genome assembly and annotation statistics.

|  | *Monodopsis* sp C73 | *Monodopsis* sp C141 | *Vischeria* sp C74 |
| --- | --- | --- | --- |
| Assembly size | 59.70 Mb | 60.47 Mb | 106.49 Mb |
| Contigs, total number | 33 | 43 | 55 |
| Contigs, with telomere | 29 | 27 | 40 |
| Contigs, telomere-telomere | 22 | 10 | 13 |
| Contig N50 | 2.24 Mb (n=11) | 2.04 Mb (n=12) | 3.09 Mb (n=14) |
| Contig N90 | 1.44 Mb (n=24) | 1.12 Mb (n=27) | 1.51 Mb (n=33) |
| Predicted protein-coding genes | 13,969 | 13,933 | 18,346 |
| BUSCO, genome assembly | 96% | 99% | 98% |
| BUSCO, predicted genes | 100% | 100% | 100% |

To gain a better picture of the genetic diversity, we generated Illumina data for two additional strains: *Monodopsis* C143 and *Vischeria* C101. SNP densities between the *Monodopsis* strains (C73, C141, and C143) ranged from 34 to 44/kb, and 10/kb between the *Vischeria* strains (C74 and C101)(Table S3). It is interesting to note that while the *Monodopsis* strains share nearly identical 18S rDNA sequences (>99.78%; Fig. S1), the genomes exhibit substantial structural and nucleotide differences (Fig. 2). This indicates that, at least in eustigmatophytes, the commonly used 18S rDNA barcode might not properly reflect the underlying genomic diversity and hence underestimate the species richness.

**Fig 2.**
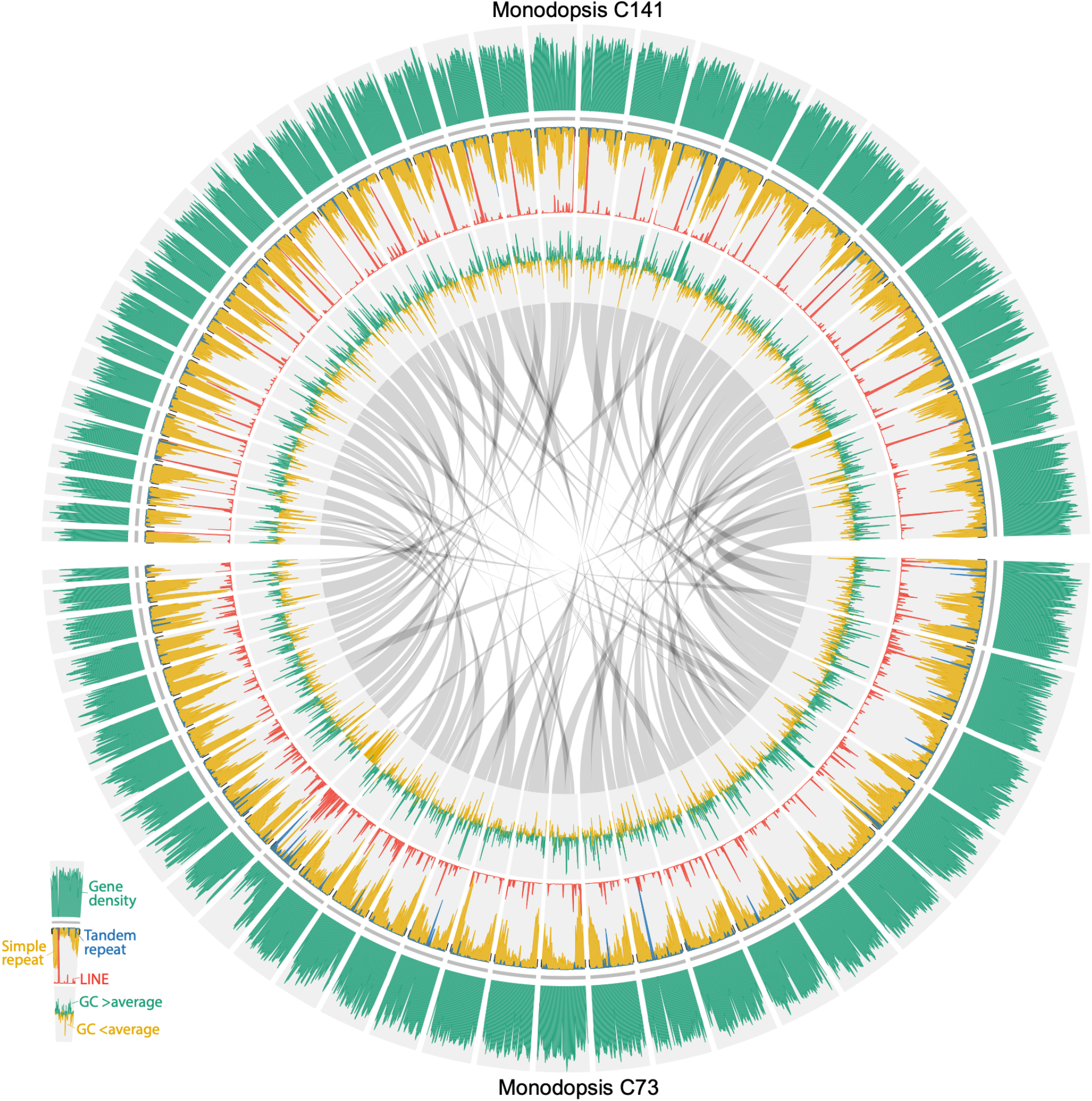
Structures of the two *Monodopsis* genomes. Simple repeats (in yellow) are particularly abundant toward the ends of chromosomes. LINEs (in red), on the other hand, tend to be locally concentrated in the middle of chromosomes (especially in *Monodopsis* C141) and likely represent centromeric regions. Extensive structural variation can be found comparing the two *Monodopsis* genomes, despite their almost identical 18S sequences. Contigs shorter than 500kb were not plotted.

### A new annotation of Nannochloropsis gaditana genome

While three *N. gaditana* genome assemblies have been published to date, two of them (B-31 and CCMP526) were based on short-read technologies and therefore had low contig N50 length (40.5Kb for B-31 and 15.3Kb for CCMP526) as well as low BUSCO completeness scores (Fig. 1A)(Radakovits et al. 2012; Corteggiani Carpinelli et al. 2014). Only the *N. gaditana* CCMP1894 genome was assembled using long reads (Schwartz et al. 2018), but unfortunately its annotation has not been published. Here we used publicly available RNA-seq data and protein evidence to annotate the *N. gaditana* CCMP1894 assembly. This new annotation has a much-improved BUSCO score (94% complete) compared to the previous *N. gaditana* annotations (40% and 85%)(Fig. 1A).

### Unusually high percentages of simple sequence repeats

*Monodopsis* and *Vischeria* have considerably larger genomes than those of *Nannochloropsis*, which can be partly attributed to their higher percentages of repetitive elements (Fig. 1B). The simple sequence repeats (SSRs) and long interspersed nuclear elements (LINEs) are particularly noteworthy. While LINEs are absent in *Nannochloropsis*, they cover around 2.9-3.6% of the *Monodopsis* and *Vischeria* genomes (Fig. 1B). SSRs have similarly expanded representations, accounting for 11.7-12.2% of the genomic content in *Monodopsis* and 20.8% in *Vischeria* (Fig. 1B). While these SSRs can be found throughout the chromosomes, they are particularly enriched toward the chromosome ends (Figs. 2 and 3). The frequencies of SSRs observed here are in fact among the highest of all genomes sequenced to date. For example, the human body louse genome (*Pediculus humanus corporis*) had the highest SSR density according to Srivastava *et al.* (2019). When reanalyzed with the same repeat annotation pipeline used here, we found SSRs account for 16.9% of the *P. humanus corporis* genome, making *Vischeria* C74 (at 20.8%) the most SSR-dense genome known to date. Future comparative studies incorporating additional genomes across eustigmatophytes are needed to clarify the impact of such high abundance of SSRs on genome structure and evolution.

**Fig 3.**
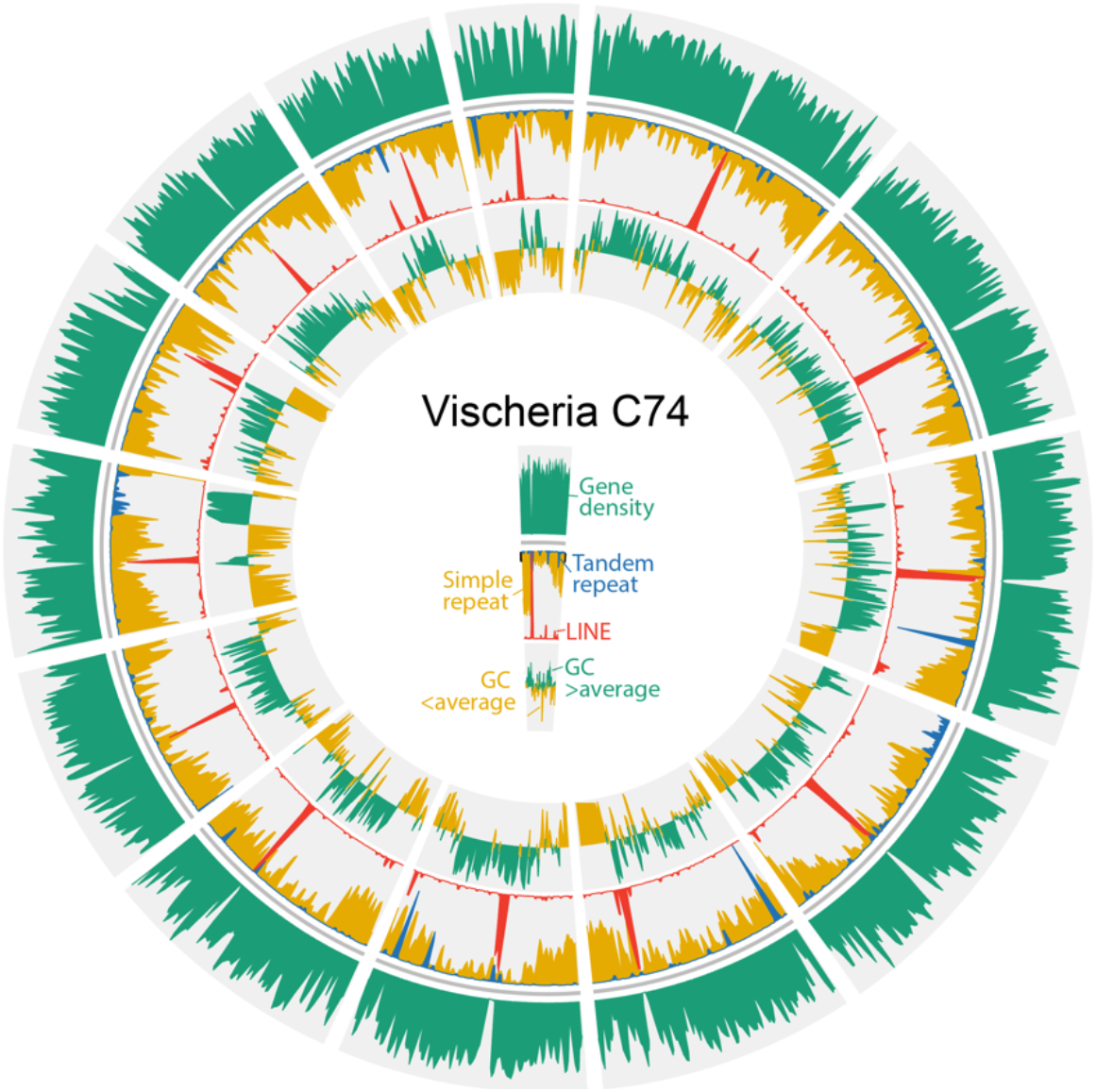
Structure of the *Vischeria* genome. Simple repeats (in yellow) are particularly abundant toward the ends of chromosomes. LINEs (in red), on the other hand, tend to be locally concentrated in the middle of chromosomes and likely represent centromeric regions. For clarity, only telomere-to-telomere contigs were plotted.

### Putative centromeric regions that are enriched in LINEs

Only a few centromere structures have been experimentally characterized in Stramenopiles. In the oomycete *Phytophthora sojae*, the centromeric regions are particularly rich in the *Copia*-like retroelements (Fang et al. 2020), whereas in the diatom *Phaeodactylum tricornutum*, the centromeres are AT-rich but devoid of repetitive elements (Diner et al. 2017). No putative centromeric region has been identified in *Nannochloropsis* to date nor in any other eustigmatophyte. Our analysis of *Monodopsis* and *Vischeria* genomes suggest that their centromeres might be characterized by islands of LINE clusters. The distributions of LINEs in *Monodopsis* and *Vischeria* are highly heterogeneous, usually with a sharp peak toward the middle of a chromosome (Figs. 2 and 3). It is likely that such LINE-dense (and gene-poor) regions function as centromeres, but further immunolabeling studies are needed. If confirmed, it would also suggest that *Nannochloropsis* might have a substantially different centromere organization given their absence of LINE.

### A haploid-dominant life cycle

The complete life cycle of eustigmatophytes has not been characterized, and no sexual reproduction has been observed. We found that several meiosis-specific genes are present in *Monodopsis* and *Vischeria*, which is consistent with what was found in *Nannochloropsis* (Table S4)(Radakovits et al. 2012; Corteggiani Carpinelli et al. 2014) and suggests eustigmatophytes do have cryptic sexual stages. In addition, we were able to identify homologs encoding flagella-related proteins in both *Monodopsis* assemblies (Table S4), despite zoospores never having been documented in *Monodopsis* (but known in *Vischeria*)(Hibberd 1981; Eliáš et al. 2017). Another missing piece of information about the life cycle of eustigmatophytes is the dominant ploidy level. While earlier genomic studies on *Nannochloropsis* suggested they are monoploid (Pan et al. 2011), no information is available for other members of eustigmatophytes. In order to assess if there is any heterozygosity present in our *Monodopsis* and *Vischeria* strains, we mapped Illumina reads to the respective genomes. We found very few SNPs could be called, and the vast majority of the alternative alleles were supported by low percentages of reads (Fig. S2), suggesting these SNPs were artifacts of residual sequencing and/or assembly errors. Therefore, we infer both *Monodopsis* and *Vischeria* have a haploid-dominant life cycle similar to *Nannochloropsis*.

### Terpenoid biosynthesis pathways differ between Monodopsis/Vischeria and Nannochloropsis

Terpenoids are an important class of natural products and have high bioenergy potentials. There are two pathways for terpenoid biosynthesis: the mevalonate pathway (MVA) and the non-mevalonate pathway (MEP). Many Stramenopiles, such as diatoms, have both pathways, while all the *Nannochloropsis* genomes sequenced to date have only the MEP pathway. Interestingly, in the *Monodopsis* and *Vischeria* genomes, we were able to find intact MVA and MEP pathways present (Fig. S3). Because *Nannochloropsis* is nested within *Monodopsis* + *Vischeria*, the most likely scenario is that *Nannochloropsis* secondarily lost the MVA pathway. This finding highlights the importance of having biodiverse genomes to infer the biology of eustigmatophytes.

### Presence of spliced leader trans-splicing and operons

Our initial analysis of the RNA-seq data revealed a low read mapping rate (~85%), which is surprising given the high genome completeness and continuity. One possible explanation is the presence of spliced leader *trans*-splicing (SLTS), which was reported in *Nannochloropsis gaditana* in a patent application (Seshadri et al. 2018). SLTS is a special mRNA maturation process, in which the 5’ end of a pre-mRNA is capped by a spliced leader (SL) sequence that is transcribed from a separate SL locus. The main function of SLTS is to add the necessary 5’ caps to each cistron in a eukaryotic operon (Lasda & Blumenthal 2011). A diverse group of organisms have been shown to have SLTS, including nematodes, cnidarians, and several unrelated protist lineages (Bitar et al. 2013; Krchňáková et al. 2017).

Upon closer inspection with SL detection pipelines, we found evidence of a single SL type in *Monodopsis* and *Vischeria*, and also confirmed the SL previously reported in *Nannochloropsis gaditana* (Table 2). The main variants of these SLs were supported by at least 155,671 reads, ensuring confidence in their accuracy (Table S5). All species also possess several minor SL sequence variants at much lower read coverage (Table S5). The main SL variants were *trans*-spliced to 12,313-17,426 AG acceptor sites throughout the genomes. Between 48% and 82% of annotated genes were located within at most 100 bp of an SLTS acceptor site (Table 2), and we observed up to 11 SLTS sites per gene (Table S5). This may suggest a complex genome-wide landscape of alternative SLTS in all species, similar to kinetoplastids (Nilsson et al. 2010). The main SL variants were encoded by 24-239 candidate SL RNA genes. Except for *Monodopsis* C141, all species possess at least two dissimilar SL RNA gene variants, which may indicate the presence of pseudogenes (Table S5). Functional SL RNA copies are expected to possess a T-rich region (*Sm* binding motif) that is required for interaction with the splicing machinery (Stover et al. 2006). We found the canonical *Sm* binding motif ATTTTG (Bitar et al. 2013) in six out of 170 SL RNA genes in *Vischeria*, but not in *Monodopsis* and *Nannochloropsis* (Table S5). This may indicate that the more recently diverged species *Monodopsis* and *Nannochloropsis* have an altered SLTS machinery with different *Sm* motifs, which will require functional molecular studies to elucidate. The secondary structures of the SL RNA genes of all species display at least one major stem loop (Table S5), consistent with SL RNAs in dinoflagellates (Zhang et al. 2007) and tunicates (Ganot et al. 2004), but divergent from the typical three-loop structure in most other organism groups (Krchňáková et al. 2017).

**Table 2:**
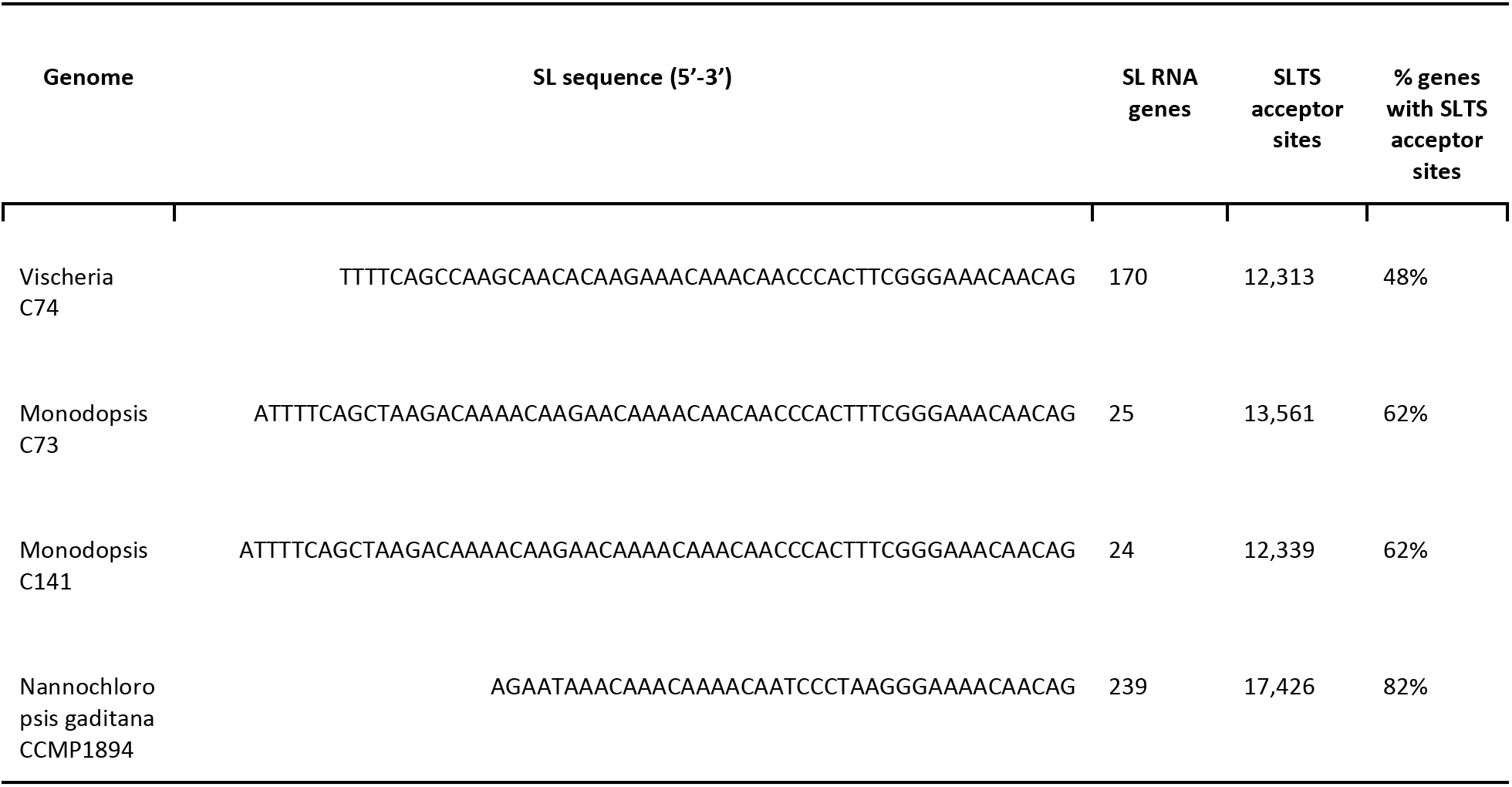
Summary of spliced leaders identified in *Monodopsis, Vischeria* and *Nannochloropsis*. The main SL sequence variant is presented with the numbers of SL RNA candidate genes, the numbers of SL trans-splice (SLTS) acceptor sites, and the percentage of genes located at most 100 bp downstream of an SLTS acceptor site. Details for SL variants and SL RNA genes are provided in Table S5.

Having established the presence of SLTS in all species, we then tested whether the physical locations of genes that receive SLs may imply the presence of operons. We first reconstructed the 5’ UTRs of gene annotations aided by the identified SLs, which yielded improved annotations for 40-80% of genes (Table S6). Using these improved annotations, we then detected SLs at 36% of genes in *Vischeria*, 58-61% in *Monodopsis* and 89% in *Nannochloropsis*. Requiring downstream genes in operons to receive the SL and intercistronic distances to be no greater than 1000bp predicted 682-1253 operons per species, containing 8-30% of all genes (Table 3). Only 21-44 of these operons had intercistronic distances of at most 100bp (Table S6). Consistent with the much higher SLTS rate, 90% of the putative *Nannochloropsis* operons receive the SL at both upstream and downstream genes, whereas *Vischeria* and *Monodopsis* show upstream STLS at only 44-64% of the putative operons. We found no significant (FDR ≤ 0.1) GO or KEGG enrichment in operonic genes compared to the full genomic background, contrary to expectations from other organisms (e.g., Zeller (2010)). This may suggest that operon evolution in these species was not necessarily driven by functional coordination of gene expression.

**Table 3:** Summary of operons predicted in *Monodopsis, Vischeria* and *Nannochloropsis* on the basis of spliced leader *trans*-splicing. Predictions required intercistronic distances of at most 1000bp and did not require SLTS at upstream operonic genes. The table presents the percentage of genes receiving SL reads, the numbers of operons, the percentage of operons where the upstream operonic gene receives SL reads, the numbers and percentages of operonic genes, and the median intercistronic distances among operonic genes. Details for SL read quantification and operon prediction using alternative criteria are provided in Table S6.

| Genome | % genes SL trans-spliced | Operons | % operons with SLTS upstream genes | Operonic genes | % total genes | Median intercistronic distance (bp) |
| --- | --- | --- | --- | --- | --- | --- |
| Vischeria C74 | 36% | 682 | 44% | 1,408 | 8% | 564 |
| Monodopsis C73 | 61% | 1,164 | 64% | 2,442 | 17% | 542 |
| Monodopsis C141 | 58% | 1,068 | 60% | 2,216 | 16% | 554 |
| Nannochloropsis gaditana CCMP1894 | 89% | 1,253 | 90% | 2,765 | 30% | 655 |

While these predictions are likely not exhaustive and will require functional validation, they are entirely consistent with other organisms where a single SL is added to both monocistronic and operonic genes, for example tunicates (Ganot et al. 2004) and platyhelminths (Boroni et al. 2018). Although SLTS has been reported in some algal lineages (Kuo et al. 2013; Roy 2017), our results provide the first insight into the genome-wide landscape of SLTS and putative operons in several eustigmatophyte algae. Future long-read RNA or cDNA sequencing will help to better define these operons and clarify the functional significance.

## Conclusion

Here we present three high-quality genome assemblies of *Monodopsis* and *Vischeria*. We found that in many aspects, *Monodopsis* and *Vischeria* genomes are substantially different from those of *Nannochloropsis*. For instance, *Monodopsis* and *Vischeria* genomes are 2-3 times larger, and boast one of the highest proportions of simple repeats among sequenced eukaryotic genomes. The centromeric regions in *Monodopsis* and *Vischeria* might be made up by LINE repeats, which are notably absent in *Nannochloropsis*. In addition, while *Nannochloropsis* lacks the MVA pathway for terpenoid biosynthesis, both MVA and MEP are present in *Monodopsis* and *Vischeria* and likely represent the ancestral state.

On the other hand, we also identified important features that are shared among the eustigmatophyte genomes. Notably, our finding and the initial characterizations of SLTS unraveled a new aspect of eustigmatophyte biology. We anticipate our new genomic data and associated analyses will greatly facilitate future research to better understand the biology of eustigmatophytes, and to better capitalize on their translational potential.

## Materials and Methods

### Data statement

The nanopore and Illumina sequencing reads were deposited in NCBI SRA under the BioProject PRJNA730568. The genome assemblies and annotations are available through NanDeSyn data portal (Gong et al. 2020) and https://figshare.com/s/c4bf156c2764ba410c30.

### Strain isolation

The three *Monodopsis* (C73, C141, and C143) and two *Vischeria* (C74 and C101) strains sequenced here were isolated from surface-sterilized bryophytes. The localities can be found in Table S1. We followed the methods outlined in Nelson *et al.* (2019) for cleaning and sterilizing the bryophyte thalli, as well as for establishing unialgal cultures that grew out from the plants. These new algal cultures are available through UTEX Culture Collection of Algae.

### Genome sequencing

We sequenced the genomic DNA on both Oxford Nanopore MinION device as well as Illumina NextSeq500 platform. Nanopore libraries were prepared using the Ligation Sequencing kit (SQK-LSK109), and sequenced on MinION R9 flowcells (FLO-MIN106D) for 60 hours or until the flowcells died. We carried out basecalling using Guppy v3.0.3 with the high accuracy flip-flop mode. For *Monodopsis* C73 and C141 strains, reads shorter than 15kb were discarded prior to assembly, and for *Vischeria* C74, a threshold of 5 kb was used. For Illumina libraries, we followed the general protocol of Nelson *et al.* (2019) using the SparQ DNA Frag & Library Prep kit and Adapter Barcode Set A. The libraries were pooled with nine other samples and sequenced on one Illumina NexSeq500 mid-output flowcell (150bp paired-end) at Cornell Institute of Biotechnology. Reads were trimmed and quality-filtered by fastp v0.20.1 (Chen et al. 2018).

### RNA sequencing

Cells grown on BG11 solution under 12/12 dark/light cycle and 22°C were harvested by centrifugation and disrupted by a SPEX SamplePrep 1600 MiniG tissue homogenizer. RNA was extracted using Sigma Spectrum Plant Total RNA kit, and strand-specific RNA-seq libraries were made by YourSeq Duet RNAseq Library Kits from Amaryllis Nucleics. The RNA libraries were pooled with 16 other samples and sequenced on one lane of Illumina NovaSeq6000 S-Prime flowcell (150bp paired-end). Reads were trimmed and quality-filtered by Trimmomatic v0.39 (Bolger et al. 2014).

### Genome assembly

We first estimated the genome size based on the K-mer frequency of Illumina reads using MaSuRCA v3.3.2 (Zimin et al. 2013, 2017). To assemble the Nanopore reads, we used Flye v2.4.1 (Kolmogorov et al. 2019) with four iterations of built-in polishing, followed by one round of medaka v0.7.1 (https://github.com/nanoporetech/medaka) processing. The nanopore assemblies were further error-corrected by Illumina reads using pilon v1.23 (Walker et al. 2014) with 4 iterations. To better assemble the telomeric regions, we used teloclip v0.0.3 (https://github.com/Adamtaranto/teloclip) to recover telomeric nanopore reads that can be aligned and appended to the contig ends. Organellar genomes were assembled separately using either GetOrganelle v1.7 (Jin et al. 2020) with Illumina reads, or Flye with a subset of nanopore reads that mapped to organellar genomes of closely related species. The Flye organellar assemblies were polished by pilon until no correction can be made. Finally, the organellar genomes were BLASTn to the nuclear genome assembly to identify and remove any redundant organellar contigs.

### Repeat annotation

Our initial repeat analysis revealed a large percentage of simple microsatellite repeats, which caused RepeatMasker to make many spurious matches to other repeat classes. To address this, we first identified and masked the simple repeats from the genome using RepeatMasker, before building the custom repeat database with RepeatModeler2 (Flynn et al. 2020). RepeatMasker was then used again to annotate and mask all the repeat classes from the genomes. Tandem repeats were identified separately using Tandem Repeats Finder (Benson 1999).

### Gene model prediction

Gene predictions were done by BRAKER2 v2.1.5 (Brůna et al. 2021), integrating both protein and transcript evidence with --etpmode and --softmasking flags on. To provide transcript evidence, we mapped RNA-seq reads to the corresponding genome using HiSAT2 v2.1.0 (Kim et al. 2015). To compile the protein evidence, we first used MAKER2 (Holt & Yandell 2011) to train SNAP (Korf 2004) on *Monodopsis* C73 based on reference-guided transcriptome assembly from Trinity v2.1.1 (Grabherr et al. 2011) and *Nannochloropsis* protein records from GenBank. The resulting gene models were then annotated with eggNOG v5.0 (Huerta-Cepas et al. 2019), and only genes with annotations were kept as the protein evidence for BRAKER gene prediction. We used the same approach to annotate *N. gaditana* CCMP1894 genome, with transcript evidence from three publicly available RNA-seq datasets (SRA accessions: SRR5152511, SRR5152512, and SRR5152516) and protein sequences from *N. gaditana* B31 and *N. salina* CCMP1776. To filter out spurious gene models from BRAKER2, we removed genes that failed to meet all of the following criteria: (1) a TPM expression level at least 0.001, (2) has functional annotation from eggNOG, and (3) was assigned into orthogroups when including all the focal eustigmatophyte genomes in an OrthoFinder v2.3.12 (Emms & Kelly 2019) run. We used BUSCO v4.0.6 (Simão et al. 2015) to assess the completeness of genome assemblies and annotations with the “Stramenopiles” lineage dataset. The final gene sets were functionally annotated (including GO and KEGG) by eggNOG v5.0. KEGG pathways were reconstructed using the KEGG Mapper tool (Kanehisa & Sato 2020).

### SNP calling

For each genome, we used bwa v0.7.17 (Li & Durbin 2009) to map Illumina reads to self as well as to the related genomes. We then use bcftools v1.9 (Li 2011) to call SNPs and keep those with quality over 50 and read depth over 20.

### Phylogenetic relationship of currently available eustigmatophyte genomes

We compiled a list of the eustigmatophyte genomes that have annotations available (Fig. 1), and used Orthofinder v2.3.12 to infer gene orthology. A total of 1,302 single-copy loci were identified, and protein sequence alignments were done by MAFFT (Katoh & Standley 2013). We then carried out phylogenetic reconstruction using IQ-TREE v2.0.3 (Nguyen et al. 2015) on the concatenated alignment matrix with automatic model selection (Kalyaanamoorthy et al. 2017) and 1,000 replicates of ultrafast bootstrapping (Hoang et al. 2018).

### Identification of spliced leader trans-splicing

We identified spliced leaders (SLs) in the C73, C74 and C141 strains as well as *Nannochloropsis gaditana* CCMP1894 (RNA-Seq library SRR10431616 from SRA) using SLIDR 1.1.4 with distance-based clustering (Wenzel et al. 2021). We relaxed the SL length limit (-x 1.25), required GT/AG splice sites and disabled the *Sm* binding motif filter. Identified SL RNA genes were inspected and aligned using MAFFT v7.407. Secondary sequence structures were inferred using RNAfold Web Server (Gruber et al. 2008). Identified SL *trans*-splice acceptor sites were compared against gene annotations using BEDTools 2.28.0 (Quinlan & Hall 2010).

We then tested whether genome-wide SL *trans*-splicing events may indicate the presence of operonic gene organisation using SLOPPR 1.1.3 (Wenzel et al. 2021). Since SLOPPR requires accurate gene annotations, particularly at the 5’ end, we first predicted 5’ UTRs guided by identified SLs using UTRme (Radío et al. 2018), relaxing maximum UTR length to 10,000 bp and maximum UTR ORF length to 400 amino acids. Reads containing at least 8bp of the SL at the 5’ end were then identified and quantified against transcript annotations using SLOPPR. Operon inference was tested with four intercistronic distance cut-offs (infinity, 1000bp, 100bp and automatic inference) and did not require upstream operonic genes to be SL *trans*-spliced. The functional annotations (GO, KEGG) of candidate operonic genes were tested for overrepresentation against the genome-wide background using hypergeometric tests in ClusterProfiler 3.14.2 (Yu et al. 2012).

## Acknowledgment

This study was supported by the National Science Foundation Dimensions of Biodiversity grant (1831428) to F.-W.L., and the Czech Science Foundation grant 20-27648S to M.E.

