## Supplementary material for "*Monodopsis* and *Vischeria* genomes elucidate the biology of eustigmatophyte algae": Fig. S1-3

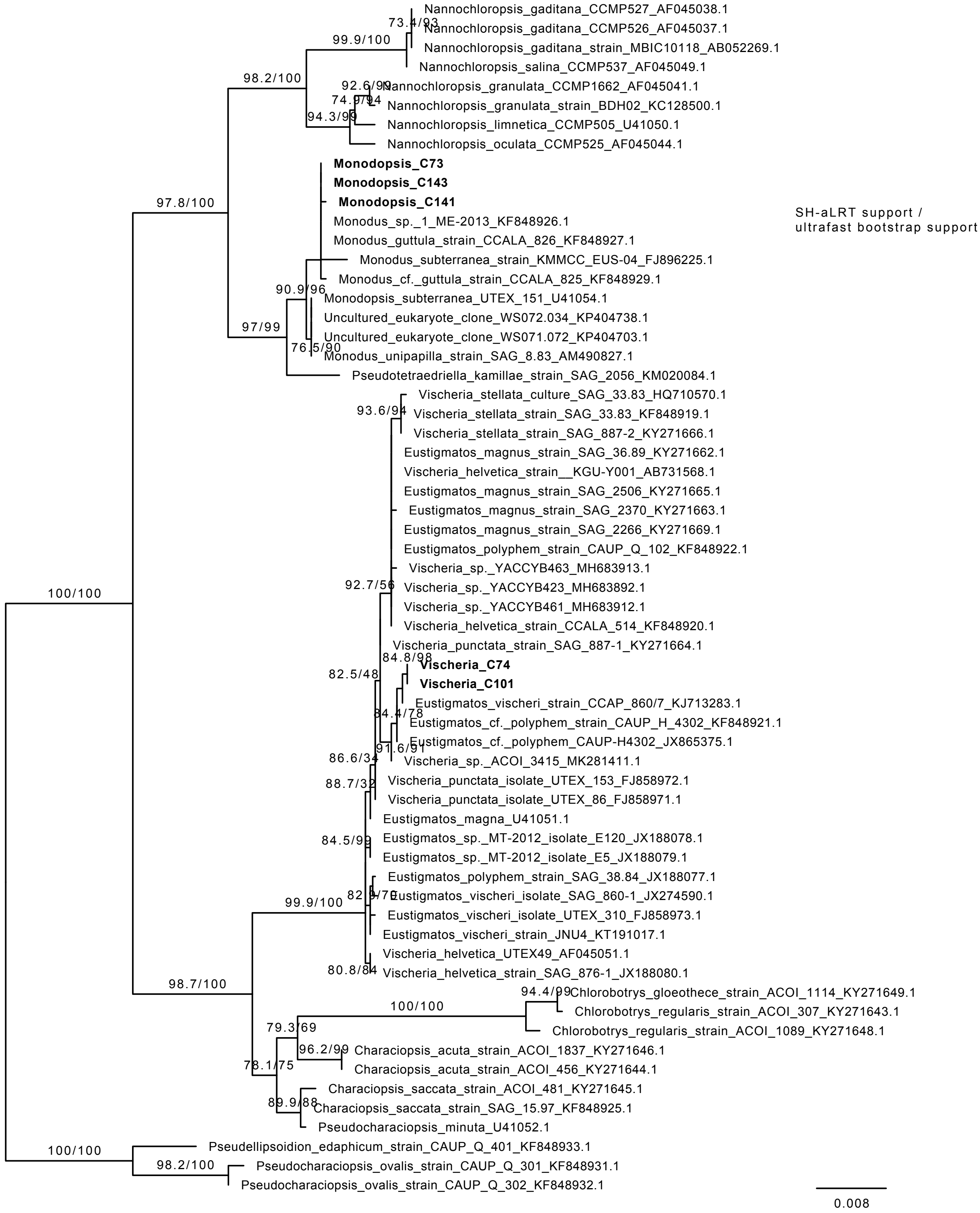


Fig. S1. **18S rDNA phylogeny of selected eustigmatophyte isolates.** The new isolates introduced in this study are in bold. The numbers above the branches are: SH-aLRT support / ultrafast bootstrap support, and are omitted if SH-aLRT support is zero.

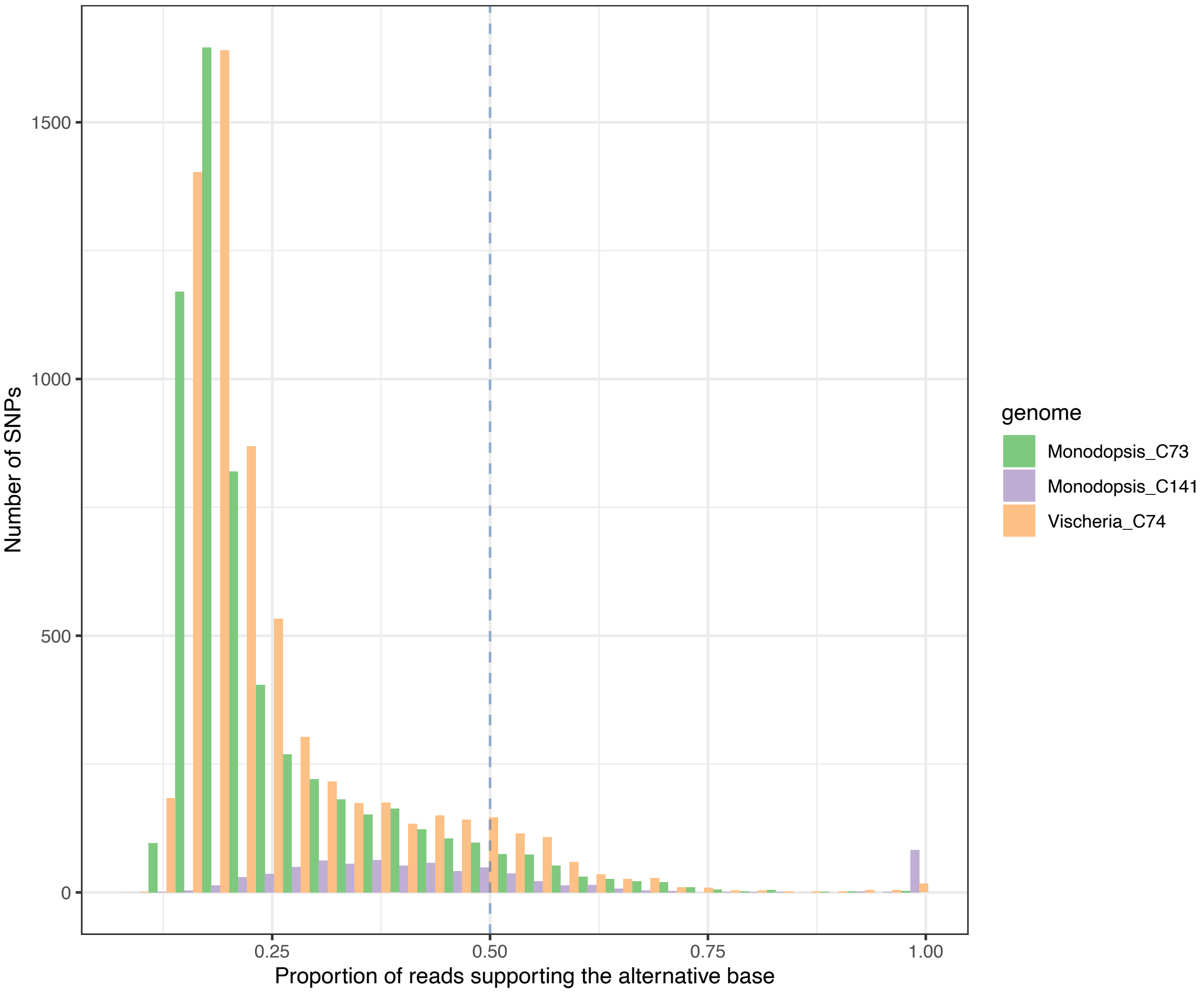


Fig. S2. **The three genomes sequenced here are likely haploid.** Very few SNPs were found while mapping Illumina reads to the respective genomes, and most of the SNPs were supported by a low percentage of reads. In other words, we found no strong evidence of heterozygosity.

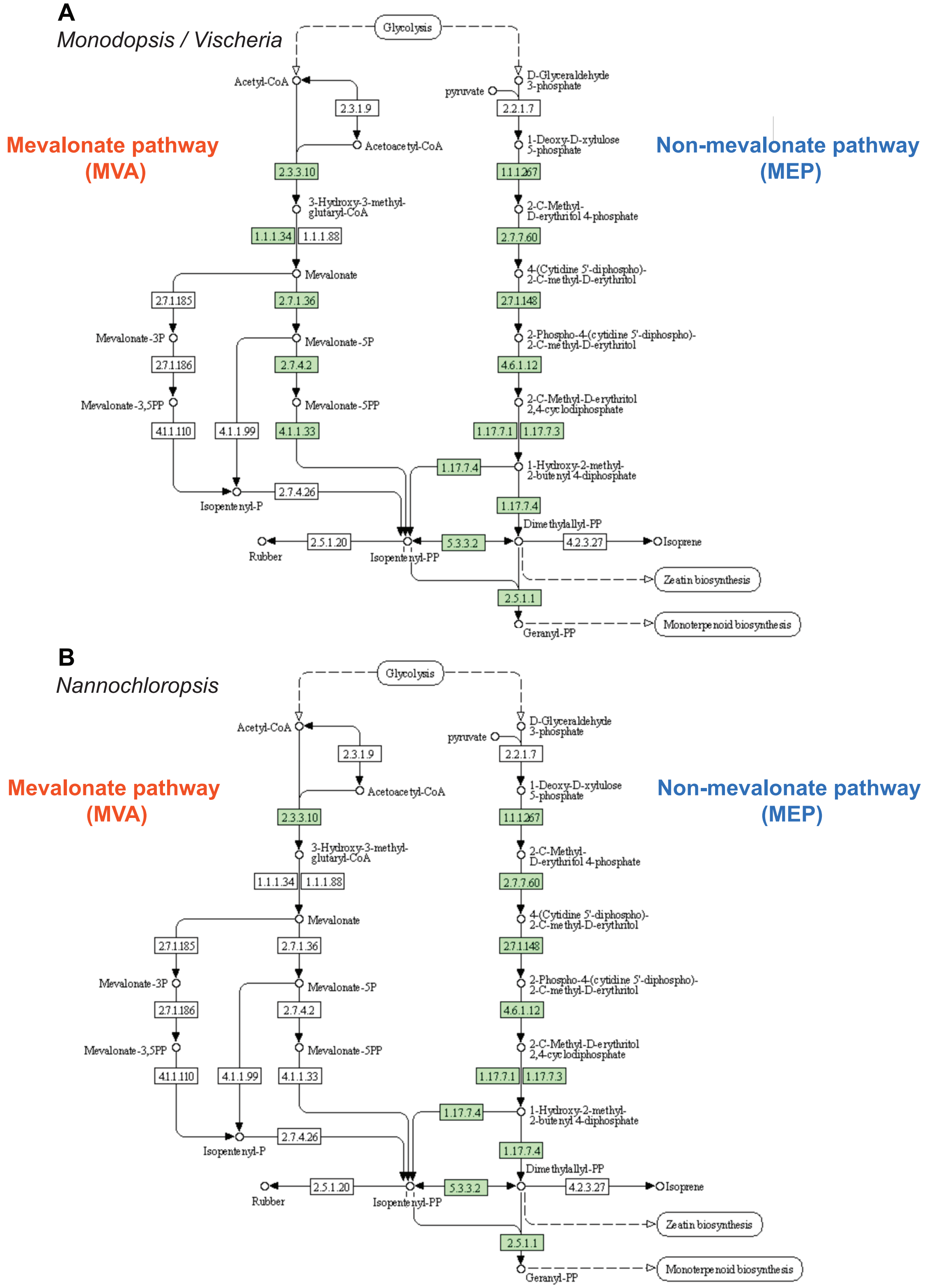


Fig. S3. **Terpenoid biosynthesis pathways.** (A) *Monodopsis* and *Vischeria* genomes have genes encoding for both mevalonate pathway (MVA) and non-mevalonate pathway (MEP). (B) *Nannochloropsis*, on the other hand, has only the MEP pathway. The pathway maps were drawn by KEGG mapper.
